# Modeling the impact of optimized airflow and sick pen management on the spread of infectious diseases in swine barns

**DOI:** 10.1101/2024.03.13.584486

**Authors:** Maryam Safari, Christian Fleming, Jason A. Galvis, Aniruddha Deka, Felipe Sanchez, Gustavo Machado, Chi-An Yeh

**Author notes:** Corresponding author: *Email address:* (Gustavo Machado).

## Abstract

The airborne spread of infectious livestock diseases plays a crucial role in the propagation of epidemics, particularly in populations confined to densely populated facilities, such as commercial swine barns. Therefore, quantitative assessments for the performance of barn ventilation systems may serve as an alternative biocontainment control strategy to reduce the spread of infectious pathogens. In this study, we present a framework to simulate airborne disease dissemination within swine barns and facilitate the strategic design of control actions, including optimization of ventilation and placement of sick animals (sick pen). This framework is based on a susceptible-infected-recovered (SIR) model that accounts for the between-pen disease spread within swine barns. A pen-to-pen contact network is used to construct a transmission matrix according to the transport of airborne respiratory pathogens across pens in the barns, via our Reynolds-averaged Navier-Stokes computational fluid dynamics (CFD) solver. By employing this CFD-augmented SIR model, we demonstrated that the location of the sick pen and the barn ventilation configuration played crucial roles in modifying disease dissemination dynamics at the barn level. In addition, we examined the effect of natural ventilation through different curtain adjustments. We observed that curtain adjustments either suppress the disease spread by an average of 56.5% or exacerbate the outbreak potential by an average of 5.7%, compared to the scenario where side curtains are not raised. Furthermore, we optimize the ventilation configuration via the selection and placement of ventilation fans through the integration of the CFD-augmented framework with the genetic algorithm to minimize the dissemination of swine disease within barns. Compared to regular barn ventilation settings, our optimized ventilation system significantly reduced disease spread by an average of 43.2%. Our study emphasizes the role of airborne transmission and a strategy for sick pen management in controlling the spread of within-barn disease.

## 1. Introduction

Although mathematical models have been used to estimate the epidemiological consequences of swine disease introduction, existing models do not often account for within-barn dissemination pathways, such as the airborne transmission (Pozzi et al., 2021; Sørensen et al., 2020; Ssematimba et al., 2022; Tuominen et al., 2022, 2023). The speed in which pathogens propagate within swine farms depends on multiple factors, including within-herd disease prevalence, pathogen’s virulence (Machado et al., 2022; Malladi et al., 2022), and population factors such as farm size, production types, barn sizes, and barn ventilation systems (Busch et al., 2021; Machado et al., 2022; Mighell & Ward, 2021). While it has been recently demonstrated the advantages of air filtration systems and the reduction of new disease introduction events (Corzo et al., 2010; Havas et al., 2023), air filters are only suitable for modern barns that can support negative pressure. However, such mechanical ventilation systems have been limited to farms in the Midwest and areas densely swine populated (Alonso et al., 2013). The most common barn ventilation system utilized in downstream farms (e.g. finisher, wean-to-finisher) remains a hybrid ventilation system that takes advantage of natural and mechanical ventilation. These barns are often called a curtain-sided facility (Godyń et al., 2020).

Of particular importance to our proposed within-barn transmission model, several studies have described the role of airflow in the within-farm dissemination of several diseases (Cador et al., 2017; Friese et al., 2012; La et al., 2021; Rosen et al., 2018; Seo et al., 2015; Vianney et al., 2023). While the disease transmission model showed significant differences in the number of newly infected pigs when modeling different levels of between-pen nose-to-nose (direct) contacts (Ssematimba et al., 2022), other models have focused on considering distance-dependent transmission within barns (Nielsen et al., 2017). The role of contaminated environment and airborne transmission showed that the combination of both pathways is sufficient to allow methicillin-resistant *Staphylococcus aureus* to persist in swine herds (Sørensen et al., 2020; Tuominen et al., 2022). However, a significant gap exists in modeling the airborne transmission explicitly via the propagation of infected droplets throughout the barn, considering the effects of natural and mechanical ventilation.

The within-barn dissemination of infectious disease triggers swine producers to either cull sick pigs or move pigs that are falling behind, sick or injured, at any time during the postweaning, growing, and finishing periods, as well as sows into an isolation pen known as a sick/recovery pen (Fraser et al., 2013). Isolating sick animals has a significant impact in curving pen-to-pen dissemination (Go et al., 2023) while also allowing sick pigs to recover without competing with healthy animals. Furthermore, to reduce within-barn disease spread, swine producers also implemented additional biosecurity measures, such as prohibiting equipment from being shared between barns and wearing personal protective equipment (disposable coveralls, gloves, and rubber boots) (Alarcón et al., 2021; Plut et al., 2023). While numerous research recommendations exist to enhance biosecurity by reducing disease introduction into barns, little is known about the effects of barn ventilation systems and the choice of sick pens on barn-level disease dissemination. There needs to be more understanding of the combined effects of ventilation and the location of sick pens that significantly hinder the effects of within-barn disease control measures. Therefore, there is an urgent need to consider such within-barn dynamics to advance our understanding of disease propagation and its implications for control strategies and decision-making.

Over the past decades, computational fluid dynamics (CFD) has become a reliable and widespread toolset to assess the influence of ventilation systems on the transmission of airborne pathogens released from bio-sources (Chen et al., 2022; La & Zhang, 2019; Lipinski et al., 2020; Motamedi et al., 2022; Yang et al., 2015). However, the combined effects of a ventilation system and sick pens on barn-level disease dynamics remain unclear. This study aims to close this knowledge gap by integrating CFD and an epidemiological compartmental model, considering the sick pen location, to predict the within-barn disease propagation. The objectives of our present study are:

1. Developing a barn-level between-pen disease dissemination model that focuses on that pathway of airborne transmission by informing the model with barn airflow;
2. Investigating the influence of sick pen location on the disease spread dynamics;
3. Investigating the influence of the barn curtain usage in curtain-sided facilities on the disease spread dynamics;
4. Exploring optimized ventilation designs that mitigate the disease spread rate via model-informed decisions for selecting and placing ventilation fans.

This simulation study is outlined as follows: We first introduce the layout of a commercial swine barn considered in this study in section 2.1. The CFD-augmented between-pen disease dissemination model developed in the current study will be presented in section 2.2. The genetic-algorithm-based optimization for ventilation design will be discussed in section 2.3. The influence of the sick pen location and curtain usage will be examined in section 3.1 and 3.2, respectively. The performance of the optimized ventilation settings in curving the disease spread will be discussed in section 3.3. We will close our discussions by providing comments on the limitations of the present approach in Section 4 and offer concluding remarks in 5.

## 2. Material and methods

### 2.1 Swine barn layout

We consider the disease dissemination within commercial-scale no-breeding barns (e.g. nursery, finishing) in a two-dimensional setting. An example barn-room is shown in Fig. 1, which accounts for the room population of 960 pigs, with an average of 30 pigs per pen. The room’s internal space is divided into several pens without imposing physical restrictions on the airflow. The pens are arranged in two longitudinal rows with a one-meter-wide walkway separating the room’s rows. The location within the barn used to keep sick animals, referred to as the sick pen or hospital pen, is also highlighted. Commercial swine barns often have ventilation fans, including intake and exhaust fans. These fans are also shown in the example barn-room. As a curtain-sided facility, the left and bottom walls of the room are covered by curtains that can be opened (rolled up) or closed (rolled down). Interior measurements of the barn-room, including the length and widths of the room and pens and the installation locations of the ventilation fans, are also shown in Fig. 1. Additionally, flow velocities of the ventilation fans in this barn were measured at the distance of 0.5 m away from the center of each fan. Note that a solid wall (top wall in Fig. 1) splits this barn into two identical rooms. However, since there is no opening in the partition wall, our study will only focus on the disease spread in one barn-room.

**Figure 1.**
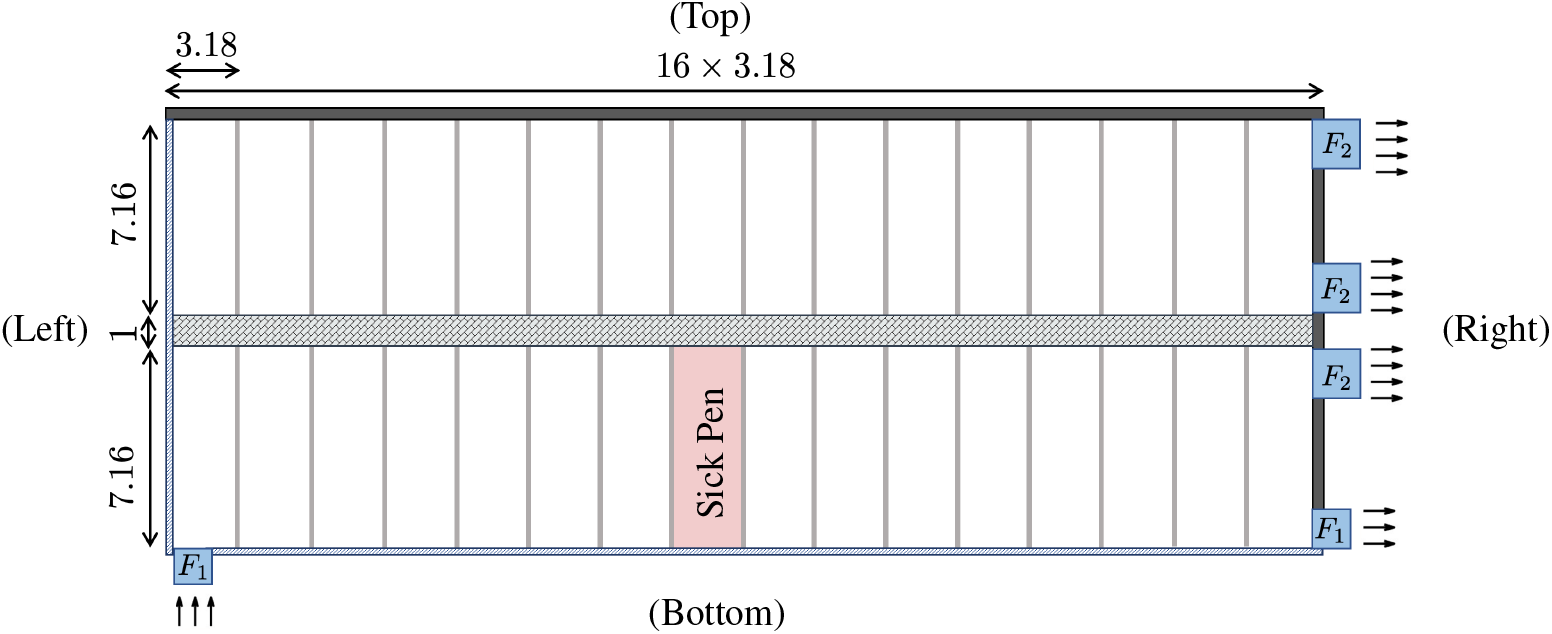
The two-dimensional model of a wean-to-finisher barn room (all dimensions are in meters). Here, *F*_1_ and *F*_2_ are ventilation fans with diameters *D*_1_ = 1.2 m and *D*_2_ = 1.45 m, velocity magnitude of *u*_fan 1_ = 2.7 m/s and *u*_fan 2_ = 3.6 m/s, respectively.

### 2.2 Barn-level disease dissemination model

#### 2.2.1. SIR model

The barn-level infectious disease model used in this study is based on the susceptible-infectious-recovered (SIR) compartmental model (Anderson & May, 1991; Diekmann & Heesterbeek, 2000; Hethcote, 2000; Newman, 2018), in which the pens are considered the epidemiological units. For each pen labeled by an index *i*, the SIR model describes the time evolution of the susceptible (*s*_*i*_), infected (*x*_*i*_), and recovered (*r*_*i*_) fractions of pigs in pen *i* by time-integrating the ordinary differential equations over 175 days of housing period at a time step Δ*t* = 1*/*5 day.

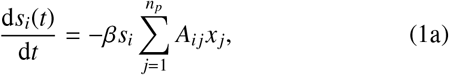

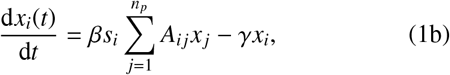

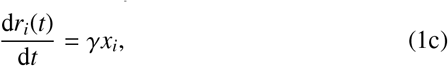

The time period of 175 days was chosen since it is the average time period for pigs to be housed in a wean-to-finisher barn (Galvis et al., 2022). Here, *n*_*p*_ is the total number of pens. The disease infection rate, *β*, quantifies the probability per unit-time for a susceptible individual to become infected. The recovery rate, *γ*, quantifies the probability per unit of time for an infected individual to recover. In this study, we choose *γ* = 0.0067/day according to the recovery period of porcine reproductive and respiratory syndrome virus (PRRSV), which can persist in pigs for an average of 150 days after the initial infection (Allende et al., 2000). For *β*, we will consider a range of values over *β* ∈ [0.2, 1] to examine its effect on inter-pen disease transmission and the optimal ventilation configuration. Most importantly, *A*_*i j*_ is the factor of disease spread from pen *j* to pen *i*. The center of our approach is to obtain a physics-based metric for the inter-pen transmission factor, *A*_*i j*_, which constitutes the *i*-*j* entry of the pen-to-pen transmission matrix ***A***.

Here, we consider that airborne transmission was the only pathway to infect pens. This transmission pathway is undertaken when airborne pathogens transport from a source pen (pen *j*) to a destination pen (pen *i*) either by the advection of the barn airflow or via self-diffusion. We rely on the CFD simulations discussed below to capture the transport phenomenon of airborne pathogens with which the airborne transmission pathway can be quantified in *A*_*i j*_.

#### 2.2.2. Computational fluid dynamics

We developed a CFD solver to simulate the airflow and the transport of airborne pathogens within a barn. The solver solves the two-dimensional, incompressible RANS equations and a passive scalar transport equation for the airborne pathogens given by

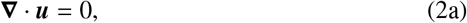

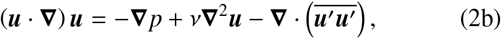

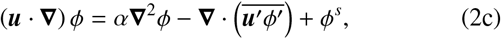

using the *k*-*ω* turbulence model (Patankar, 2018; Wilcox, 1998, 2008) (Appendix A.) Here, ***u*** is the velocity field of the airflow, *p* is the pressure field, and *ϕ* is the concentration field of the airborne pathogens. Since the airflow is of concern, *ν* is the kinematic viscosity of the air, and for *α*, we consider the diffusivity of carbon dioxide in the air as a general airborne substance. The source term in the transport equation, *ϕ*^*s*^, denotes the generation rate of the airborne pathogens due to the respiratory droplets released by sick pigs located in a specific pen. This in-house solver has been validated against commercial CFD software packages, e.g. Ansys Fluent (Doumbia et al., 2021; Li et al., 2017; Matsson, 2023; Yeo et al., 2019).

The CFD simulation provides the barn airflow velocity field, ***u***(ξ), and the concentration field of the airborne pathogens, *ϕ*(ξ), where ξ is the space variable. While the airflow in a barn can vary against many factors, in this study, we will only focus on those associated with ventilation configurations in natural (curtains) or forced (fans) settings. Therefore, for a given barn room, we can informally express the airflow velocity field obtained from the CFD simulation as

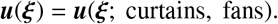

Implementing these ventilation settings (fans and curtains) in the CFD solver is discussed in Section 2.2.4. Moreover, from equation (2c), we see that the concentration field of airborne pathogen depends on the ventilation airflow built up in the barn (***u***) and the location of sick pigs (*ϕ*^*s*^). Hence, the pathogen concentration field obtained from CFD can also be denoted in short as

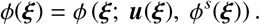

Next, we discuss how we rely on the pathogens concentration field *ϕ* subject to a pen source *ϕ*^*s*^ to quantify the pen-to-pen transmission factor *A*_*i j*_.

#### 2.2.3 Construction of pen-to-pen transmission matrix

Given a pen infection source (*ϕ*^*s*^) with sick pigs, the CFD simulation provides the concentration field of the airborne pathogens *ϕ* for all pens in the barn. In other words, given a source pen (pen *j*) where the sick pigs generate airborne pathogens, we determine the amount of airborne pathogens propagated from the source pen (pen *j*) to a destination pen (pen *i*) using the CFD simulation. Therefore, we compute each *i*-*j* entry of the transmission matrix ***A***, or *A*_*i j*_, using the following procedure:

1. Form a source term with a unit-strength generation rate uniformly distributed in the source pen (pen *j*). That is,

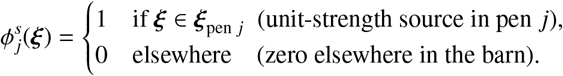

This step is demonstrated in Fig. 2 (a), where pen 1 is chosen as the source pen and the source term 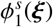 is visualized. This source term 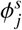 will be fed into the right-hand-side of the equation (2c) to perform the CFD simulation.
2. Run the CFD simulation to obtain the concentration field of airborne pathogens with the prescribed source from Step 1. That is, obtain from CFD

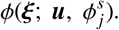

This step is demonstrated in Fig. 2 (b), where the resulting pathogen concentration field, 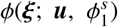, due to the source formed in Step 1 is visualized.

**Figure 2.**
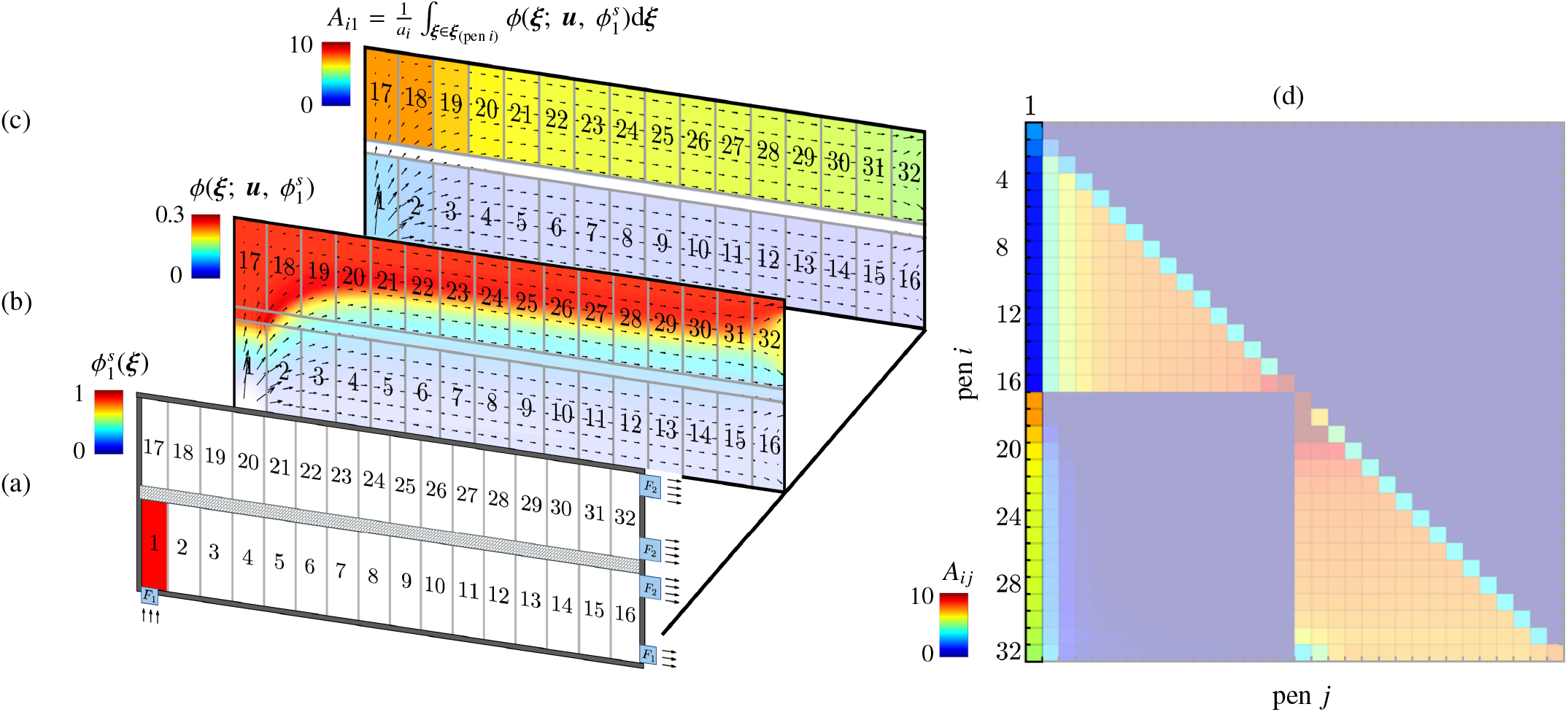
The procedure of constructing the transmission matrix ***A***: (a) Specify a unit-strength source of airborne pathogens 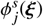 uniformly distributed in a selected source pen (pen 1 in this example); (b) Perform CFD simulation to obtain the pathogen concentration field, 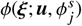, due to the source specified in pen *j*; (c) Calculate the amount airborne pathogen accumulated in each pen for *A*_*i j*_ using equation (3); (d) Each CFD run will render a full column of the transmission matrix ***A***. In this example, where the source is placed in pen 1 ( *j* = 1), column *A*_*i*1_ is obtained.
3. Calculate the total amount of airborne pathogens accumulated in each pen by spatially integrating the pathogen concentration field over the area occupied by the pen. The amount of airborne pathogens propagated to a destination pen (pen *i*) from the prescribed source pen (pen *j*) constitutes the *i*-*j* entry of the transmission matrix ***A***, or

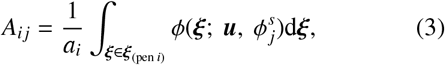

where *a*_*i*_ is the area of pen *i*. This step is demonstrated in Fig. 2 (c) by visualizing the total amount of airborne pathogens accumulated in each pen. Once a source term is formed for a prescribed source pen (the *j*-entry) in Step 1, the accumulated pathogen in all pens (all *i*-entries) will be determined from the CFD simulation in Step 2. Therefore, each CFD simulation in Step 2 will render a whole column of the transmission matrix ***A***, as shown in Fig. 2 (d).
4. Iterate Step 1 to Step 3 while prescribing the source pen to the rest of the pens in the barn until the transmission matrix ( ***A***) is entirely constructed column-by-column.

By introducing the entirely constructed transmission matrix into the SIR equations (1), the CFD-augmented disease dissemination model is ready to be used for our investigation.

Note that the use of the unit-strength concentration level of the airborne pathogen obtained from the CFD, i.e. *ϕ*(ξ; ***u***, *ϕ*^*s*^), is linearly proportional to the strength of the source *ϕ*^*s*^. This is due to the linear nature of the transport equation (2c) with respect to *ϕ* and leads to a linear proportionality between *A*_*i j*_ obtained from equation (3) and the strength of the source *ϕ*^*s*^. Recall that the source term *ϕ*^*s*^ stands for the generation rate of the airborne pathogen due to the respiratory droplets released by sick pigs per unit area. This generation rate is expected to be linearly proportional to the infected fraction *x* _*j*_ in a source pen *j*, or 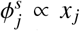 Hence, by computing *A*_*i j*_ using a unit-strength source, the transmission due to an actual source strength that varies with *x* _*j*_ is automatically accounted for by the term,

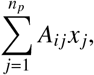

that appears in the SIR equations (1). The actual generation rate can vary against a few other factors, including the population density and the size/weight of the pigs in a source pen. While the influences of these factors are not in the scope of the present study, they can also be easily included by exploiting the linear proportionality between *A*_*i j*_ and the source strength. Our model allows for re-scaling each *A*_*i j*_ entry according to those factors (e.g., different population densities among pens), instead of performing another series of CFD simulations to construct a new transmission matrix. This feature offers promising extendability for our CFD-augmented SIR model in future studies.

#### 2.2.4 Modeling ventilation set-up through boundary condition

As noted, the concentration of the airborne pathogen depends on the airflow built up in the barn. The airflow velocity field is, in turn, dependent on the ventilation settings. Our CFD simulations model these ventilation settings by prescribing different boundary conditions. These boundary conditions and the associated ventilation settings are summarized in Table 1. Inlet/outlet velocity boundary conditions with uniform velocity profiles model the intake/exhaust fans. We assume that intake fans do not introduce additional airborne pathogens into the barn. Also, open (rolled-up) curtains are modeled by a traction-free boundary condition that allows the flow to enter or exit the domain without restrictions. In contrast, closed (rolled-down) curtains were treated in the same manner as for solid walls.

**Table 1:** The boundary conditions (BCs) used to model physical ventilation settings.

| Type | Velocity | Pressure | Concentration |
| --- | --- | --- | --- |
| Wall | $\mathbf{u} = 0$ | $\hat{\mathbf{e}}_n \cdot \nabla p = 0$ | $\hat{\mathbf{e}}_n \cdot \nabla \phi = 0$ |
| Intake fan | $\mathbf{u} = u_{\text{fan}}(+\hat{\mathbf{e}}_n)$ | $\hat{\mathbf{e}}_n \cdot \nabla p = 0$ | $\phi = 0$ |
| Exhaust fan | $\mathbf{u} = u_{\text{fan}}(-\hat{\mathbf{e}}_n)$ | $\hat{\mathbf{e}}_n \cdot \nabla p = 0$ | $\hat{\mathbf{e}}_n \cdot \nabla \phi = 0$ |
| Opened (rolled up) curtain | $\hat{\mathbf{e}}_n \cdot \nabla \mathbf{u} = 0$ | $\hat{\mathbf{e}}_n \cdot \nabla p = 0$ | $\hat{\mathbf{e}}_n \cdot \nabla \phi = 0$ |
| Closed (rolled down) curtain | $\mathbf{u} = 0$ | $\hat{\mathbf{e}}_n \cdot \nabla p = 0$ | $\hat{\mathbf{e}}_n \cdot \nabla \phi = 0$ |

Note: $\hat{\mathbf{e}}_n$ is the unit normal vector directing into the barn domain, and $u_{\text{fan}}$ is the fan speed that can vary for different fans. An open curtain refers to one that is rolled up and closed for one that is rolled down.

### 2.3 Ventilation optimization

Disease transmission via airborne pathogens depends on the concentration field that forms according to the airflow in the barn. Since barn ventilation is a critical factor on which the airflow depends, the infection curve can be flattened by an informed design of barn ventilation. Here, we seek an optimal ventilation configuration that effectively flattens the infection curve. Given a pre-selected sick pen (as shown in Fig. 1), our ventilation optimization focuses on four design variables:

1. The number of ventilation fans, or *n*_*f*_ ;

2. The speeds and diameters of fans, given by the specifications available in a list of industrial fans;

3. Fans configuration as intake and/or exhaust;

4. Fans location on the barn boundaries, except for the partitioning wall (top).

For item 1, we considered a range of 3 to 8 fans for *n*_*f*_ . For item 2, a library with information about fans from the BESS Laboratory at the University of Illinois Urbana-Champaign was considered, where 650 agricultural fan models are included with their specifications on speeds and diameters^1^. For Item 4, we also introduced a restriction that no fans are placed within 10 meters from either side of the sick pen. This restriction was imposed to reflect an industrial practice in swine barns, where sick pigs are usually kept warm, so their exposure to high-speed airflows is discouraged.

The ventilation optimization was conducted using the genetic algorithm (Satheesan, 2023; Satheesan et al., 2023; Tsang et al., 2023; Zhou, 2007) integrated with the CFD-augmented SIR model. The genetic algorithm explored 9, 000 combinations of the design variables listed above and sought the set of design variables that minimizes the cost function given by

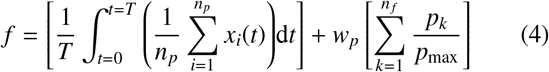

where 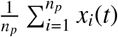 is the pen-averaged infected fraction obtained from the CFD-augmented SIR model with *β* = 1, *T* is the housing period of the pigs (175 days), *p*_*k*_ is the power consumption of fan *k, p*_max_ is the maximum power consumption by ventilation fans that is determined based on the maximum number of the fans and the power usage of the most powerful fan in the library (3 kw), and *w*_*p*_ is the penalty coefficient for the second term. Note that the first term, as the main objective in optimization, uses the average of an infected fraction over 175 days to measure the flatness of the infection curve. The second term, on the other hand, penalizes the objective function by the total power consumption of the fans to discourage the optimization to always choose the most powerful fans for excessively high ventilation rates (Kurnitski et al., 2021). In the present study, we chose *w*_*p*_ = 0.2 to emphasize the flatness of the infection curve, but its value can be tuned for other considerations in practice. The effectiveness of the optimal ventilation setting in reducing disease dissemination will be discussed in section 3.3.

## 3. Results and discussions

### 3.1. Effects of sick pen location on disease spread

The barn airflow is modeled as a constant control variable while the sick pen location is examined. We focus on the scenario where all curtains in the barn shown in Fig. 1 are closed such that the barn airflow is only driven by ventilation fans. As indicated by the vector fields in Fig. 3 (a, b), the airflow under this ventilation configuration enters the barn from the intake fan near the bottom-left corner, moves towards the top wall, gradually turns right, and forms a horizontal flow pattern until it approaches the exhaust fans on the right wall. We examined how this airflow builds different pathogen concentration fields from different source pens, and its influence on the disease dissemination in the barn.

First we rely on CFD simulations to examine how an airborne pathogen disseminates over the barn from a source pen, even before the disease dissemination between pens occurs due to the airborne pathogen. The airborne pathogen is generated only from the selected source pen in such a setting. Still, the concentration field formed by the airflow heavily depends on the location of the chosen source pen. Indeed, we observed that the formation of the pathogen concentration field depends on not only the location of the source pen in the barn but also the proximity of the source pen to the intake or exhaust fans. This is shown in Fig. 3 (a) and (b), where pen 1 and pen 31 are chosen as the source pens, respectively. Note that pen 1 resides immediately next to the intake fan, and pen 31 is close to the exhaust fans. When pen 1 is selected as the source pen, as shown in Fig. 3 (a), the airflow results in high levels of pathogen concentration over pens 17 to 32 that are located along the top wall. For the same case, the pens along the bottom boundary exhibits low levels of pathogen concentration, compared to those along the top wall. On the other hand, in Fig. 3 (b), we observed that the pathogen generated in pen 31 only accumulates in the downstream pen 32 and is immediately removed from the barn by the exhaust fans. These results demonstrate the importance of the source pen location to the pathogen concentration field that forms in the barn.

Evaluating the concentration fields formed due to different source pens is the central idea of the construction of the transmission matrix, as discussed in section 2.2.3. Therefore, we examined the transmission matrix to broaden our understanding of the influence of the source pen location beyond the two representative cases discussed in Fig. 3 (a) and (b). First, we correlate the observations from the two representative cases to those presented by the transmission matrix, as shown in Fig. 3 (c). The 1^st^ column of the transmission matrix ( *j* = 1), obtained by prescribing pen 1 as the source pen, exhibits high transmission levels over the *i*-entries from 17 to 32. This observation is closely related to the high pathogen levels spreading over the pens 17 to 32, as observed in Fig. 3 (a). On the other hand, the 31^th^ column indicates that airborne pathogens sourced from pen 31 only transmits to pen 32, as the pathogens immediately leaves the barn after passing through pen 32. Similar observations and interpretations to the 31^th^ column can be made for the 15^th^ column, where the pathogens sourced from pen 15 are immediately removed from the barn after passing through pen 16. A notable case is the 17^th^ column of the transmission matrix, where the airborne pathogens are sourced from pen 17. In this pen, a low-speed airflow is locally formed near the top-left corner of the barn. It results in high levels of accumulation of airborne pathogens in pens 17 and 18, as indicated by the associated entries of the highest level over the entire transmission matrix. Moreover, we identify a general pattern in the transmission matrix: the entries below the main diagonal exhibit higher levels of pathogen transmission than those above the main diagonal. This pattern is formed due to the nearly uniform airflow that moves from the left boundary of the barn towards the right boundary where exhaust fans are installed.

**Figure 3.**
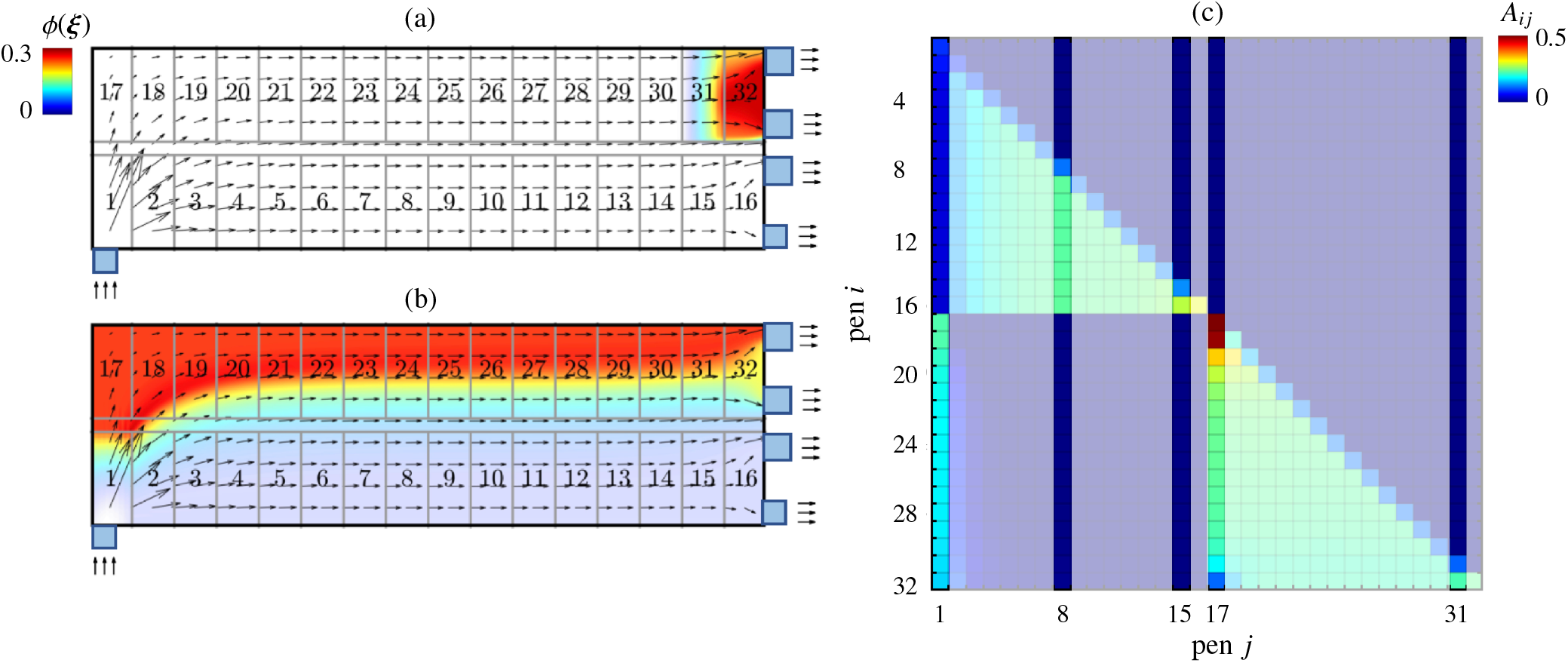
Concentration field of airborne pathogens across the barn where (a) pen 1, and (b) pen 31 are prescribed as the source pen, respectively. (c) Normalized pen-to-pen transmission matrix of regular ventilation setting where all curtains in the barn are closed.

This nearly uniform airflow dictates the upstream-downstream relationships between a source-destination pen pair, giving rise to the pattern observed in the transmission matrix. These observations once again confirm the airflow’s crucial roles and the source pen’s location for their influences on the propagation of airborne pathogens in swine barns.

Here, we utilized between-pen disease dissemination using the SIR model. Initially, we select a pen as the only sick pen in the barn, prescribe a 100%-fraction of infection to the selected sick pen, and time-integrate the SIR model with the constructed transmission matrix in Fig. 3 (c). That is, we initialize the SIR states of each pen at *t* = 0 by

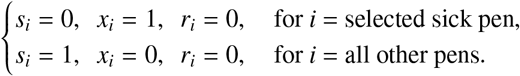

and start the time integration of the SIR model from the above initial condition, to examine how the location on the selected sick pen influences the disease spread.

We observed that an informed selection of sick-pen locations can significantly postpone the disease outbreak in the barn. This is demonstrated by comparing Fig. 4 (a) to (b), where we track the between-pen disease dissemination initiated respectively from pen 8 and pen 31 over 175 days. Note that the selection of pen 8 as the sick pen is the regular setting shown in Fig. 1, and pen 31 is located close to the exhaust fans. Here, we focus on the infected fraction, *x*(*t*), for all pens and the resulting concentration fields of the airborne pathogens in time. Day 0 is associated with the initiation of disease spread, which can be understood as the initial placement of sick pigs into the selected sick pen. Of note, the airborne pathogens are sourced from the sick pen only. Moving forward from the initial conditions, disease dissemination occurs, and infection occurs in other pens, as shown in the day 73 column in Fig. 4. The infected pens also become the sources of the airborne pathogen, significantly increasing the pathogen concentration inside the barn. This is particularly clear for the case where pen 8 is selected as the sick pen. Remarkably, when pen 31 is chosen as the sick pen, even on day 120 the infection is limited to the pens near the right boundary, as shown in Fig. 4 (b). As opposed to that case, the regular sick pen selection (pen 8) results in a pen-averaged infected fraction of over 60% on day 73, as shown in Fig. 4 (a).

**Figure 4.**
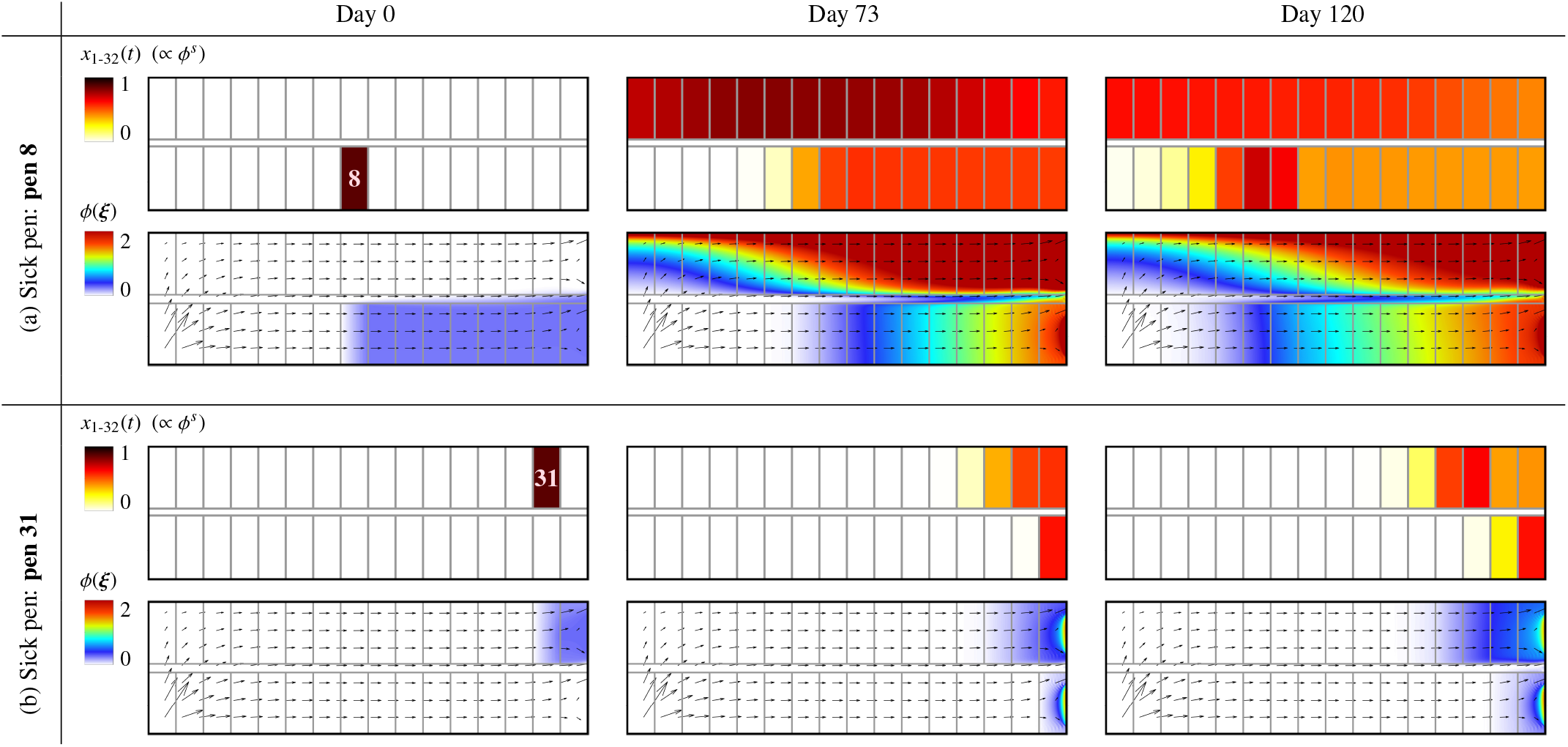
The influence of the sick-pen location on the time evolution of the infected fractions for all pens and the concentration fields of airborne pathogens: (a) pen 8 is selected as the sick pen; (b) pen 31 as the sick pen. Representative snapshots were taken on day 0 (initial condition), day 73, and day 120, post-seeded infections in the respective pens.

We further analyzed the time-resolved history of the pen-averaged infected fraction for its dependence on the sick pen location. Fig. 5 (a) presents more convincing evidence towards the effectiveness of an informed selection of sick pens in postponing the disease outbreak. When selecting the pen 1 as a sick pen, the infection peak appears only 15 days after disease dissemination is initiated. Such a rapid increase in infection is attributed to the proximity of the pen 1 to the intake fan, which promotes the dissemination of airborne pathogens from the sick pen and leads to the rapid outbreak. For the regular sick pen setting (pen 8 as the sick pen), the infection peak appears on day 73, reaching an average of 60%. When pen 31 is chosen as the sick pen, the infected fraction remains below a remarkable 10% average over the first 120 days. Although the infection increases after 130 days, the flattened infection curve over the first 120 days would allow for enough reaction time to introduce outbreak preventive measures, such as relocating sick animals to the sick pen from other pens or culling these animals. Such reduction in disease spread via the selection of sick pen holds over a wide range of infection rates. This is shown in Fig. 5 (b), where we consider the infected fractions averaged over the 175-day housing period with the infection rate, *β*, ranging from 0.2 to 1. We observed that the selection of pen 31 as the sick pen achieves lower averaged infection than the regular setting (pen 8) over the range of *β* considered. Here, we notice that the informed sick-pen selection not only reduces the total number of infected animals in the barn, but also significantly delays the outbreak and provides enough time for the producers taking measures to control disease dissemination.

**Figure 5.**
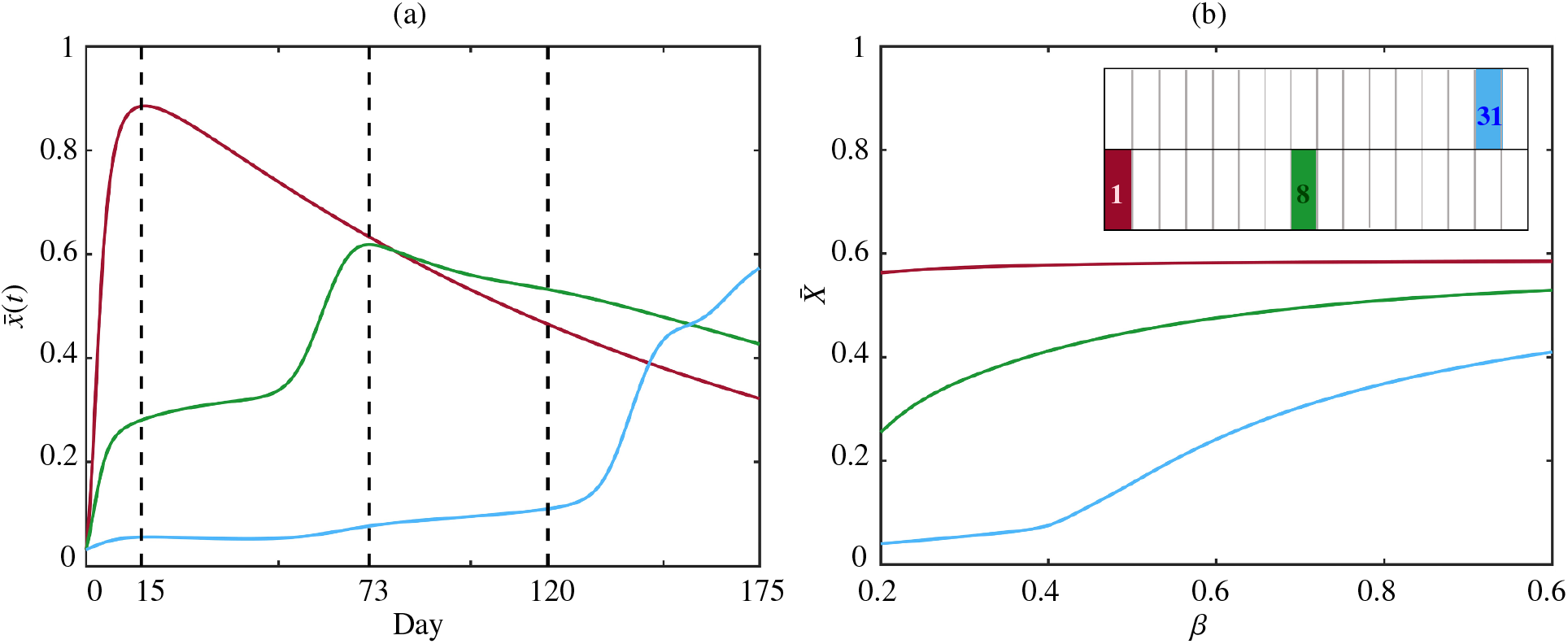
(a) Average fraction of the infected pigs over wean-to-finisher time and pens, 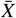, versus disease infection rate *β* for different choice of sick pens. (b) Average fraction of the infected pigs over pens, 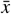, versus day for the different choice of sick pens with a disease infection rate of *β* = 0.5, where 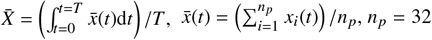, pen and *T* = 175 day.

### 3.2 Effects of curtains settings on disease spread

In this section, we study the influence of the curtain adjustments on the barn-level disease spread dynamics while holding the sick pen location and the deployment of ventilation fans as constant control variables. We focus on the regular setting in Fig. 1, where pen 8 is chosen as the sick pen, and all ventilation fans are put in action. This curtain-sided facility has curtains installed along the left and bottom boundaries, and the adjustments of these curtains are modeled through the boundary conditions discussed in Section 2.2.4. Different adjustments will build up different barn airflow, which modifies the pathogen concentration field and hence the disease spread dynamics between pens. Here, we toggle between the opened/closed setting for each curtain and consider the four scenarios:

1. both curtains are closed;
2. only the bottom curtain is opened;
3. only the left curtain is opened;
4. both curtains are opened.

First examine the airflow and the pathogen concentration fields built up by these four scenarios of curtain adjustments using CFD simulations, when considering the regular sick pen (pen 8) as the only source pen. For scenarios 1 and 3, we observed a nearly uniform left-to-right airflow built up in the barn, as shown in Fig. 6 (a) and (c). Driven by such airflow, the airborne pathogens are generally transmitted from the source pen (pen 8) to the downstream pens on its right. Opening the curtain at the bottom wall (scenario 2), as shown in Fig. 6 (b), induces a small upward flow towards the top wall over the left side of the barn and increases the overall left-to-right airflow speed than that in scenarios 1 and 3 (as indicated by the longer arrows). This higher airflow speed reduces the pathogen concentration sourced from pen 8, and the slight upward flow also slightly curves the transport trajectories of airborne pathogens. Notably in Fig. 6 (d) where both curtains are opened, we observed that the airborne pathogens sourced from pen 8 are almost immediately removed from the barn through the bottom curtain, significantly reducing the pathogen concentration. These results show clear evidence that the transmission of airborne pathogens in the barn can be modified considerably via curtain adjustments.

**Figure 6.**
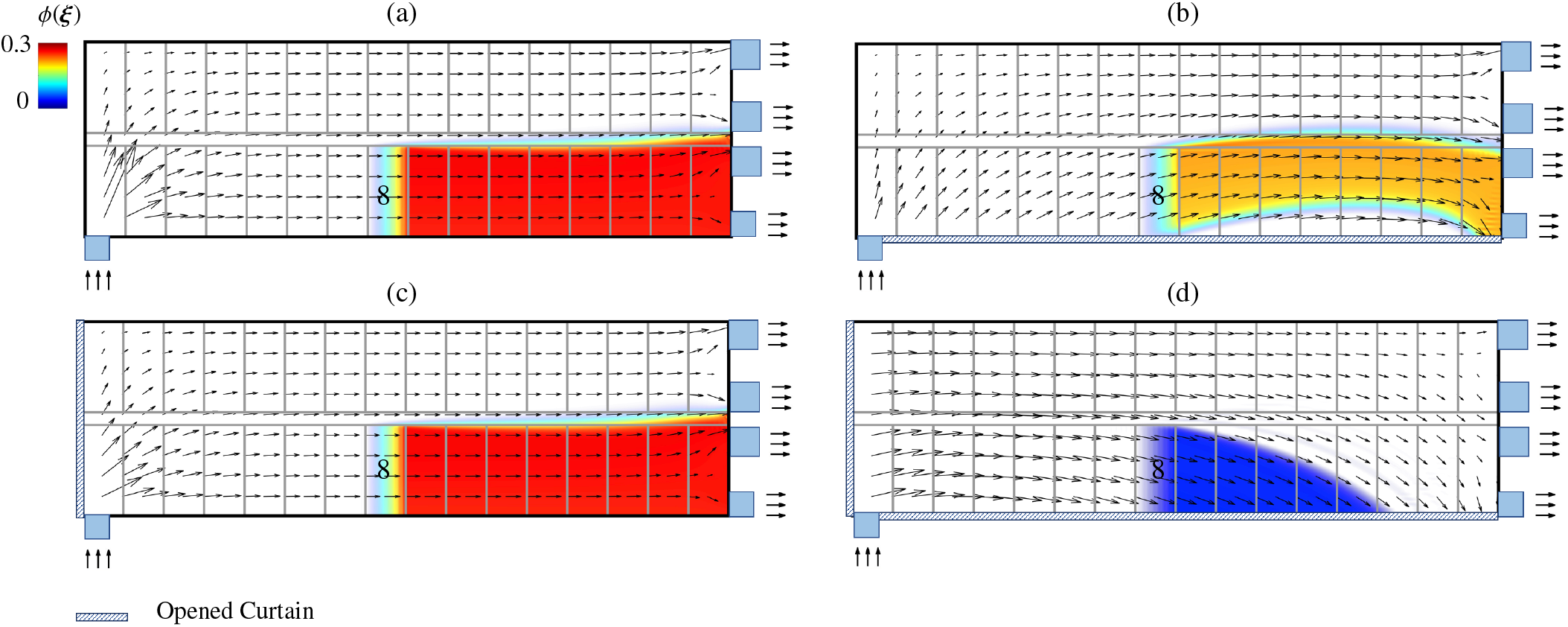
Concentration field of airborne pathogens within the barn where pen 8 is prescribed as the source pen considering four scenarios where (a) both curtains are closed, (b) only the bottom curtain is opened, (c) only the left curtain is opened, and (d) both curtains are opened.

Transmission matrices in Fig. 7 provide detail insights into the heavy dependency of the airborne pathogen transmissions on the curtain adjustments, when assessing other pens as infection source pens. We observed that in the scenario where both curtains are raised, the barn-wide transmission of airborne pathogens is significantly reduced, compared to other three curtain settings. This can be seen by comparing the transmission matrix in Fig. 7 (d) to those in (a-c). The general pattern that the entries below the main diagonal have higher magnitudes than those above the diagonal holds for the transmission matrices in Fig. 7 (a-c) (respectively associated with scenarios 1-3). This is due to the general left-to-right airflow with a slight upward velocity component locally built up near the intake fan. However, in Fig. 7 (d), where both curtains are opened, we observed that the entries above the main diagonal also have similar levels of magnitudes to those below the main diagonal. This is attributed to the barn-wide downward airflow induced by the opened bottom curtain, as seen in Fig. 6 (d). Therefore, in the scenarios where both curtains are opened, the opened bottom curtain greatly assists in the removal of airborne pathogens from the barn in addition to the exhaust fans, significantly reducing the pathogens level accumulated in the barn.

**Figure 7.**
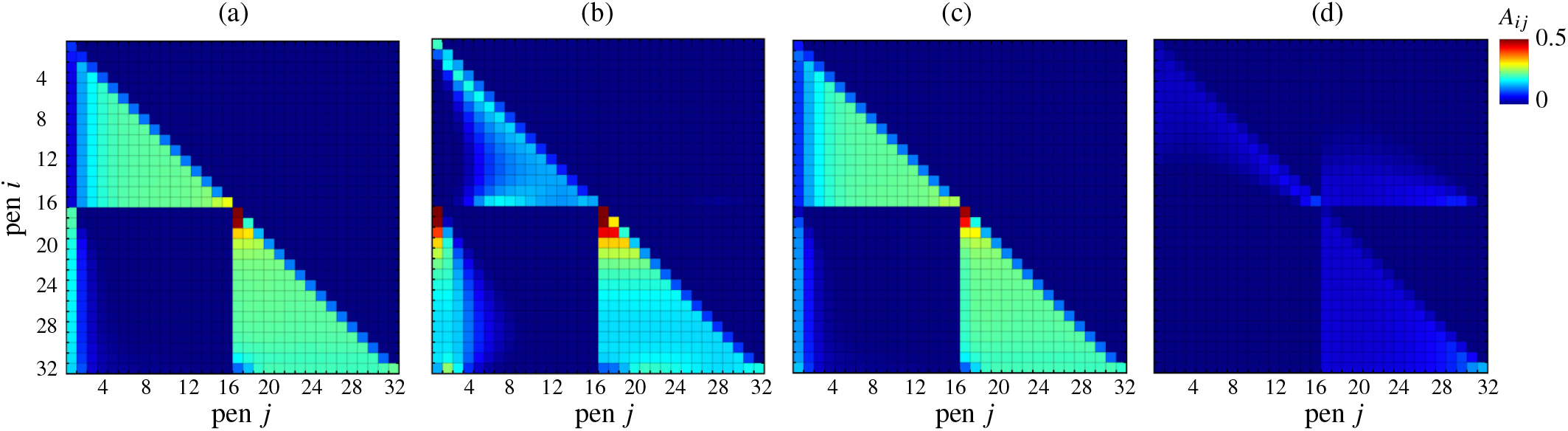
Normalized pen-to-pen transmission based on the maximum concentration of airborne pathogens considering four scenarios where (a) both curtains are closed, (b) only the bottom curtain is opened, (c) only the left curtain is opened, and (d) both curtains are opened.

Next, we initiate the disease dissemination from the regular sick pen 8 using the SIR model and the four transmission matrices constructed in Fig. 7. We observed that, with both curtains opened, the infected fractions are significantly reduced over the housing period, as shown in Fig. 8 (a). As indicated in Fig. 8 (b), this trend holds over a wide range of infection rates. A surprising observation is that opening curtain setup does not always guarantee lower infection levels: in Fig. 8, we observed that the case with opened bottom curtain exhibits higher levels of average infection than the case where both curtains are closed. This trend holds over a wide range of *β*, considering the average infection fraction over the entire housing period shown in Fig. 8 (b). This surprising outcome can be attributed to the airflow when only the bottom curtain is opened, where we observed higher upward flow speed than in other cases, as discussed in Fig.6 (b). This upward velocity enhances transmission between the pens along the bottom boundary and those along the top wall, leading to a higher infection fraction than other cases. In particular, we notice the first column of the transmission matrix in Fig. 7 (b) exhibits higher magnitudes than the first columns of the other three transmission matrices in the exact figure. This indicates that, when the airborne pathogen is sourced from pen 1, higher levels of pathogen is accumulated in the barn when only the bottom curtain is open, compared to other curtain adjustments. Fig. 9 affirms this by comparing the pathogen concentration fields for cases where no curtains are opened to the cases where only the bottom curtain is opened. We observed that the upward airflow induced by the opened bottom curtain pushes the airborne pathogen against the top wall. The airborne pathogen, while advecting towards the right boundary, remains close to the top wall such that only the exhaust fan on the top-right corner can effectively remove the pathogen from the barn, resulting in higher levels of pathogen concentration, compared to the case where both curtains are closed. In summary, in this section, we showed that the spread of barn-level disease depends on curtain adjustments, and opening curtains does not always lead to lower infection rates (Chantziaras et al., 2020).

**Figure 8.**
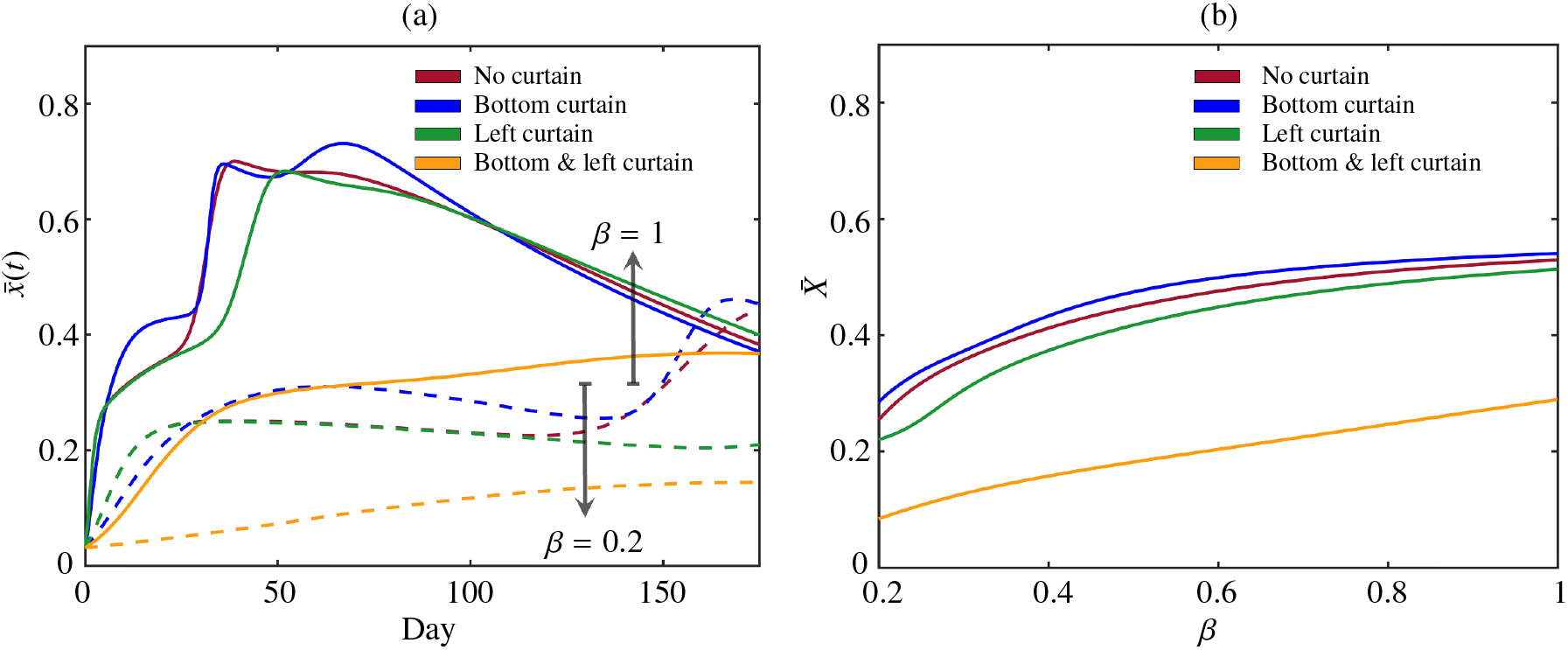
(a) The average fraction of infected pigs across all pens, 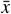, versus wean-to-finisher time for disease infection rate of *β* = 0.2 and *β* = 1, and (b) Average fraction of infected pigs over wean-to-finisher time and all pens, 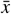, versus disease infection rate *β*, where 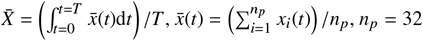 pen and *T* = 175 day.

**Figure 9.**
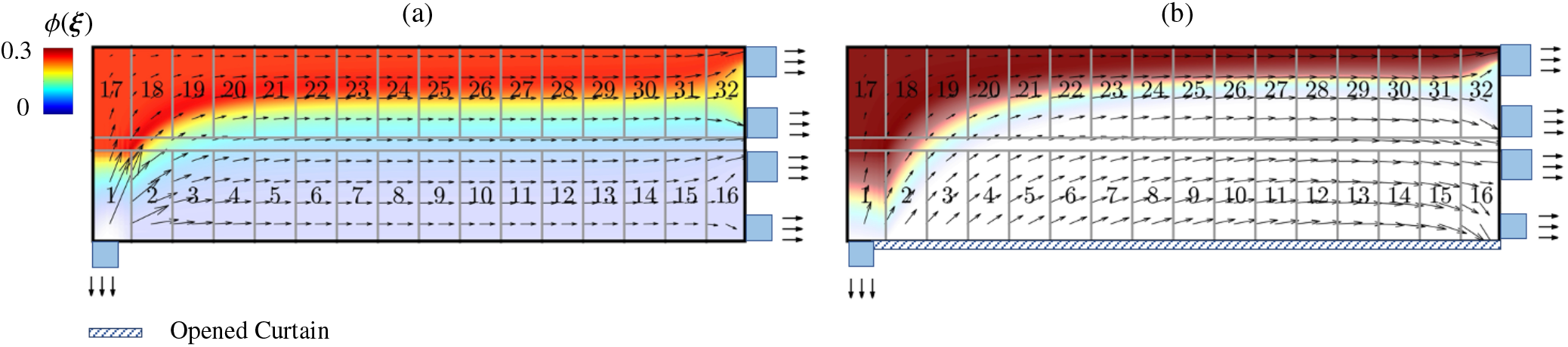
Concentration field of airborne pathogens within the barn, considering pen 1 prescribed as the source pen, where (a) both curtains are closed, (b) only the bottom curtain is opened.

### 3.3 Optimization of barn ventilation

The ventilation optimization with genetic algorithm identifies the optimal combination of design variables including number of fans, types of the fans, and their locations on the barn boundaries. Here, we assumed no opened curtains and pen 8 as the regular sick pen. Also, the optimization was conducted using the highest infection rate considered in this study, i.e. *β* = 1. Illustrated in Fig. 10 (a) is the identified optimal ventilation setting: it consists of two intake fans installed on the right boundary, an exhaust fan on the left boundary close to the aisle, and another exhaust fan on the bottom wall 10 m away from the sick pen. The airflow enters the barn from the intake fans on the left boundary and leaves through the exhaust fans on the left side of the barn, forming a nearly uniform right-to-left airflow, as shown in Fig. 10 (b). We also noticed a locally low-speed region formed near the bottom-left corner of the barn. When airborne pathogens are only sourced from pen 8, they are removed from the barn solely by the closest exhaust fan on the bottom boundary.

As opposed to the transmission matrices we present in previous sections, in the transmission matrix for the optimized ventilation we observed that the entries above the main diagonal exhibit higher transmission magnitudes than to those below the main diagonals, as shown in Fig. 10. This pattern is formed due to the nearly uniform airflow that moves from the intake fans on the right wall toward the exhaust fan on the left side of the barn. We also noticed that large-magnitude entries appear in the first four columns of the matrix. This is attributed to the local low-speed region near the bottom-left corner, resulting in the high accumulation of airborne pathogens in these pens when the pathogens are sourced from pens 1 to 3. The pattern of the 5^th^ to 16^th^ columns of the transmission matrix shows that the airborne pathogens, when being sourced from pens 5 to 16, are removed from the barn by the exhaust fan placed on the bottom wall before they spread across the pens 1 to 4. This results in low levels of pathogens concentration in pens 1-4. In contrast, the airborne pathogens sourced in the pens adjacent to the top wall, pens 20-32, leave the barn through both exhaust fans. The downward velocity component near the exhaust fan on the bottom boundary leads to the transmission of airborne pathogens from these pens to the pens 1 to 4.

**Figure 10.**
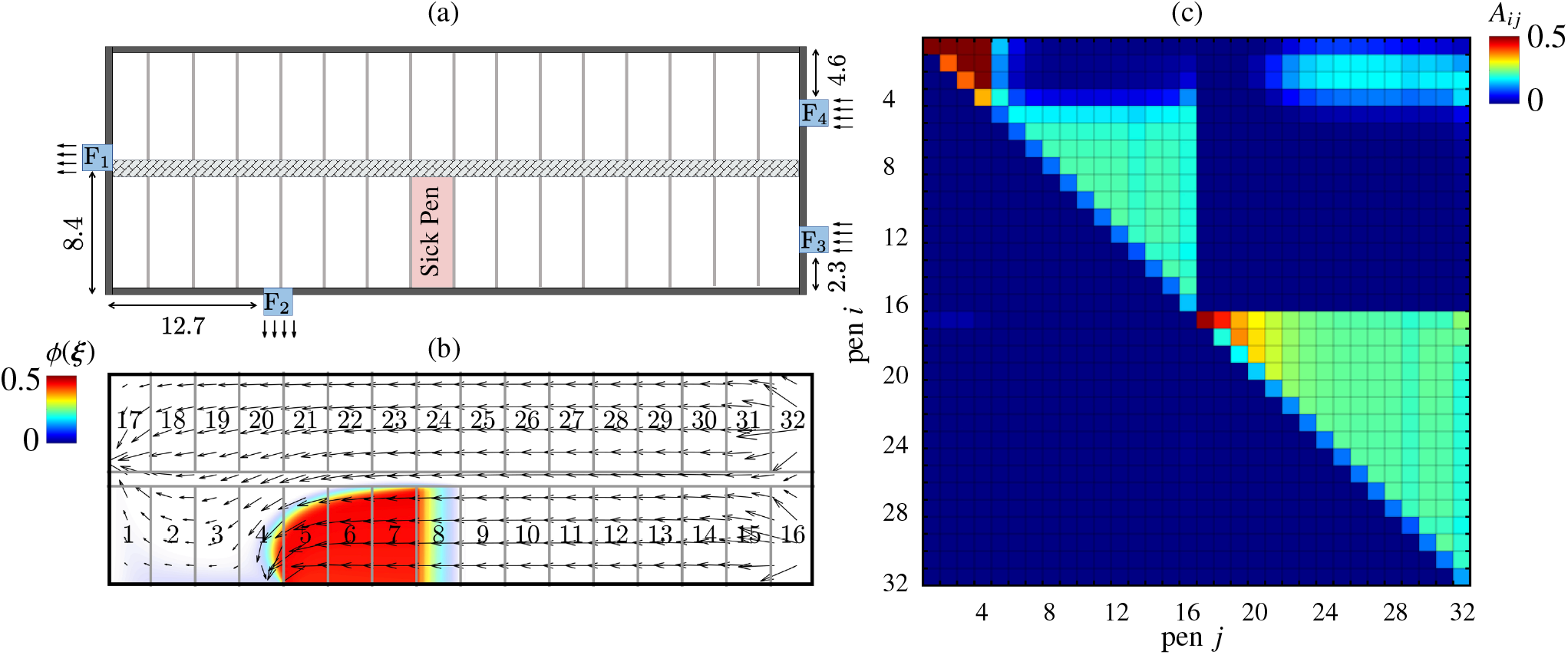
(a) Optimized ventilation setting of the swine barn where *F*_1_, *F*_2_, *F*_3_ and *F*_4_ are ventilation fans with diameter of *D*_1_ = 1.22 m, *D*_2_ = 1.22 m, *D*_3_ = 1.22 m, *D*_4_ = 1.145 m and velocity magnitude of *u*_1_ = 6.27 m/s, *u*_2_ = 6.75 m/s, *u*_3_ = 7.07 m/s and *u*_4_ = 6.34 m/s (all dimensions are in meter) . (b) The concentration of the airborne pathogens for the optimized ventilation where pen 8 is prescribed as the source pen. (c) Normalized pen-to-pen transmission for the optimized ventilation setting.

We demonstrated the effectiveness of the optimized ventilation in mitigating disease dissemination in the barn. Here, the disease dissemination is initiated from the regular sick pen (pen 8) using the CFD-augmented SIR model and the transmission matrix constructed in Fig. 10 (c). We observed that optimized ventilation effectively postpones disease outbreak with a low-ered peak value of infected fraction, compared to the regular setting. This is shown in Fig. 11, where the infected fraction in the barn with optimized ventilation is compared to that with the regular ventilation setting. At the same infection rate *β* = 1 used for the optimization, similar initial infection growth rates are observed for both cases over the first 3 days. After that, the infected fraction from the optimized setting slows down, significantly reducing the pen-averaged infected fraction. Using the same optimized ventilation setting obtained from *β* = 1, its effectiveness in suppressing disease spread also holds for lower values of the infection rate, *β*, as shown in Fig. 11 (a) with *β* = 0.2 as a representative case. Overall, the optimized ventilation reduces the infected fraction by an average of 43.2%, compared to the regular ventilation setting, over the entire range of infection rates considered, as shown in Fig. 11 (b).

Next, we assess the optimized ventilation’s robustness in suppressing disease spread against the uncertainties associated with the source pens from which the disease dissemination is initiated. When clinical signs are identified in a swine barn, many infected animals may be left undiagnosed and remain in healthy pens (Amass & Baysinger, 2006) rather than being relocated to the sick pen. In such a scenario, the barn-level disease dissemination may initiate from multiple pens in the barn on top of the sick pen instead of initiating from the sick pen only. To associate this situation with our SIR model, the undiagnosed animals located in pens other than the sick pen translate to an uncertainty in the initial SIR state from which the time integration for the SIR model starts. Here, we aim to characterize the performance of optimal ventilation in suppressing disease spread while accounting for the uncertainties and delays in disease detection. In addition to the sick pen, we randomly select many pens with an initial infected fraction averaging 15%. This fraction of 15% is chosen based on assuming that 3 to 5 infected pigs are left unidentified in a healthy pen that typically holds 25 to 35 pigs per pen. We considered 50 sets of randomly selected pens and initialized the SIR states by

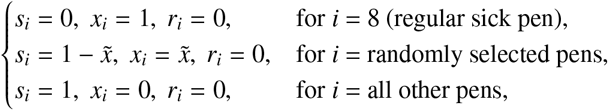

where the averaged 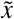 over all randomly selected pens is 15%. We started time integrating the SIR model from these 50 initial conditions.

**Figure 11.**
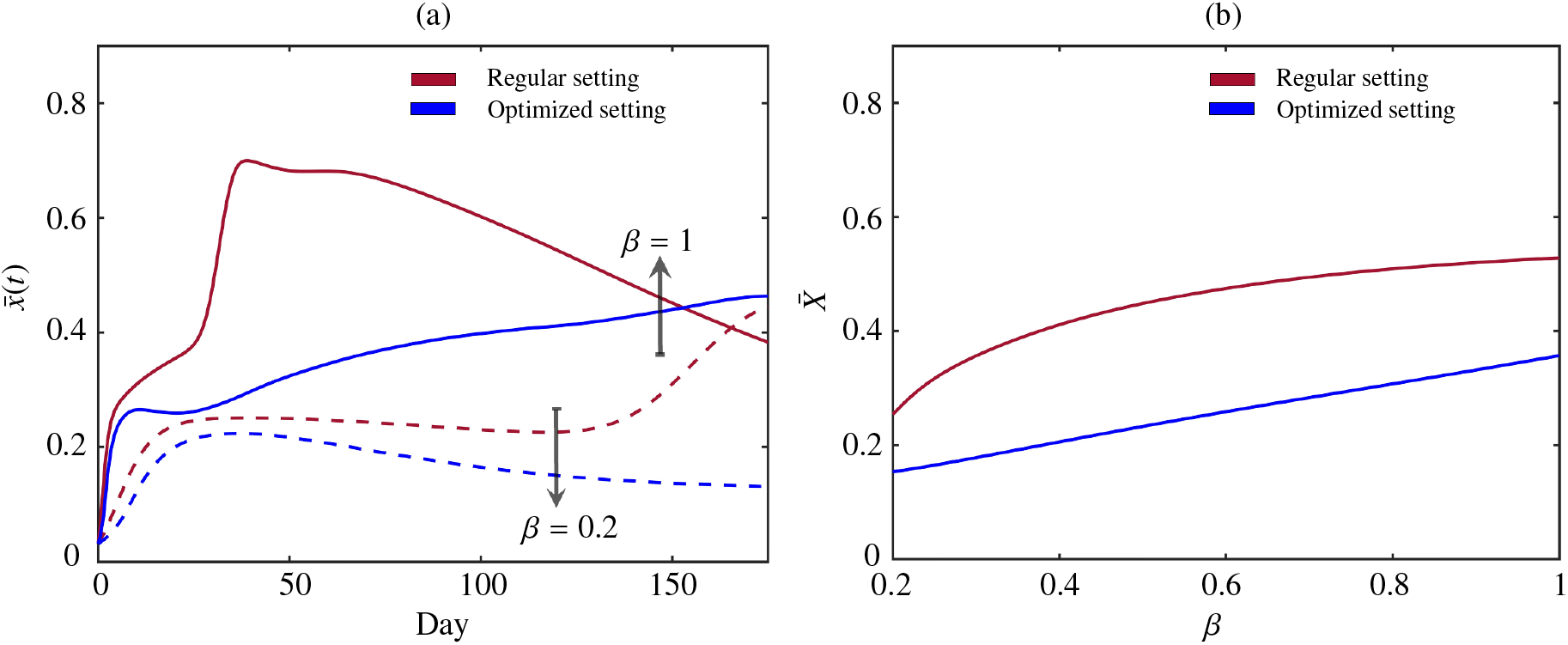
(a) Average fraction of infected pigs across all pens 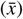 versus wean-to-finisher time for *β* = 0.2 and *β* = 1, and (b) average fraction of infected pigs over wean-to-finisher time and all pens 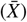 versus disease infection rate *β*, where 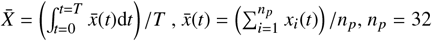 pen and *T* = 175 day.

Our results show that the effectiveness of the optimized ventilation in suppressing barn-level disease spread is robust against the source-pen uncertainties. This is demonstrated in Fig. 12, where the average infection from the optimized ventilation setting is compared to that from the regular setting for these randomly chosen initial conditions. The results from one representative initial SIR state is shown in Fig. 12 (a), where we observed that the optimized ventilation setting consistently yields a lower level of infection than the regular ventilation setting, across all infection rates considered. The same comparison between the optimized and regular ventilation setting for all 50 initial conditions is presented in Fig. 12 (b), where the highest infection rate, *β* = 1, is considered. These results show the effectiveness of the optimized ventilation in containing the barn-level disease outbreak, even when sick animals are not immediately identified and relocated to a selected sick pen due to the incubation of disease.

**Figure 12.**
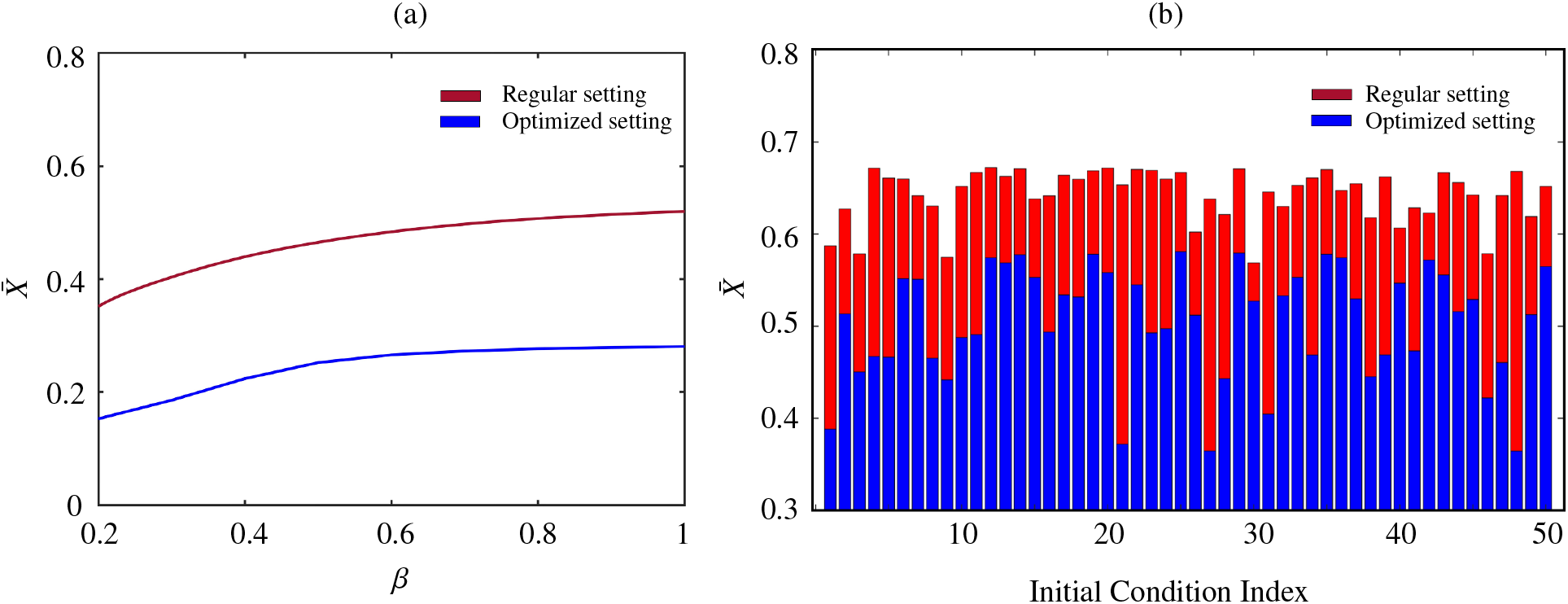
(a) Average fraction of infected pigs over time and pens versus disease infection rate *β* for a random initial condition of barn-level disease dissemination. (b) Average fraction of infected pigs over time and pens for fifty random initial conditions at disease infection rate of *β* = 1, where 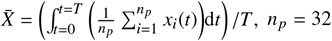 pen, *T* = 175 day.

## 4. Limitations and further remarks

We note that the present study performed the CFD simulations in a two-dimensional setting. The CFD-augmented disease dissemination model can benefit from using threedimensional CFD to obtain more realistic representations of airflow and airborne pathogens in a barn room. Another limitation of the present work is the insufficient airborne disease spread data. This includes the diffusivity of the airborne pathogens and the infection rate via airborne pathways (Andraud et al., 2013; Arruda et al., 2019; Björnham et al., 2020; La et al., 2022; Olesen et al., 2017; Seo et al., 2012). Although a range of infection rates is considered, precise knowledge of the infection rate and its dependence on the concentration of the airborne pathogen are required to accurately predict airborne disease dissemination within the barn. Study trials collecting such data for pathogens such as PRRSV could enhance the accuracy of the proposed model in predicting disease dissemination within swine production facilities. Ultimately, our optimized ventilation approach can consider the favorable environmental conditions for pigs within the barn, such as temperature, temperate fluctuation, humidity levels, and gases, which have been documented to affect the swine health and growth performance (Huynh et al., 2005; Kim et al., 2005; Vande Pol et al., 2016). Future approaches must incorporate these elements to ensure a comprehensive and effective ventilation system that fully supports the well-being and development of pigs.

## 5. Conclusions

We presented a framework to simulate the disease spread dynamics via airborne pathways in swine barns. This framework is based on the SIR model that accounts for the disease spread between pens due to the transport of airborne respiratory pathogens by the barn airflow. We employed CFD to simulate the airflow and the transport of airborne pathogens across the pens in a barn to construct the transmission matrix in the SIR model. This framework shows the crucial role of the ventilation configuration and the sick pen location in mitigating the spread of disease within the barn. We also investigated the effect of natural ventilation through different curtain adjustments. Our results showed that the curtain adjustments in the example barn can either suppress the disease spread by an average of 56.5% or exacerbate the spread by an average of 5.7%, compared to the scenario where side curtains are not raised. We demonstrate that the average fraction of infected pigs during the outbreak decreased from 58% to 22% by housing sick animals in an appropriate pen. Also, we used the genetic algorithm and the CFD-augmented model to design an optimal ventilation configuration that minimizes the outbreak probability in barns. We showed that optimized ventilation can mitigate disease transmission by 43.2% compared to regular ventilation settings. Overall, the proposed framework can effectively curve disease dissemination within swine barns, paving the way for further advancements in disease control strategies.

## Authors contribution

MS, JAG, CY, and GM conceived the study. CF and FS coordinated the data collection. MS conducted data processing and cleaning and designed and wrote the model code. MS and CY designed the computational analysis. MS, CY, and GM wrote and edited the manuscript. All the authors discussed the results and critically reviewed the manuscript. GM secured the funding.

## Acknowledgments

The authors want to acknowledge participating companies and veterinarians for their insightful comments and engaging discussions.

## Ethics Statement

The authors confirm the journal’s ethical policies, as noted on the journal’s author guidelines page. As this work did not involve animal sampling or questionnaire data collection by the researchers, ethics permits were not needed.

## Appendix

Our in-house CFD solver employs the *k*-*ω* turbulence model to solve the RANS equations (2). The the Reynolds stress term 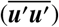 in the momentum equation (2b) is modeled by

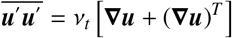

Where

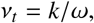

is the turbulent eddy viscosity, *k* is the turbulence kinetic energy (TKE), and *ω* is the specific dissipation rate. To close the RANS equations (2) and the two additional unknowns, *k* and *ω*, the *k*-*ω* model solves the transport equations for *k* and *ω* given by

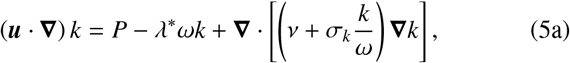

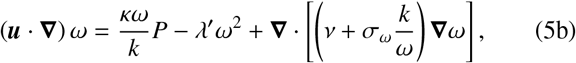

where 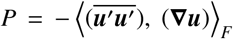 is the production of turbulence kinetic energy. Other constants in equation (5) are chosen to be *λ*^*^ = 0.09, *σ*_*k*_ = *σ*_*ω*_ = 0.5, *κ* = 0.52 and *λ*^′^ = 3*/*40 (Matsson, 2023; Patankar, 2018; Wilcox, 1998).

Boundary conditions for *k* and *ω* are also required to perform the RANS simulation. These boundary conditions vary for different ventilation settings as well. For intake fans, the boundary conditions for TKE and dissipation rate are prescribed as

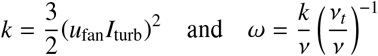

where the turbulent intensity *I*_turb_ ≡ |***u***^′^|*/u*_fan_ = 0.05 and the viscosity ratio *ν*_*t*_*/ν* = 10 are used in this study. On walls and closed curtains (Menter, 1993), TKE and dissipation rate are set to

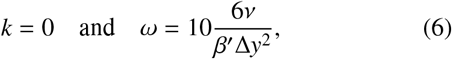

where Δ*y* is the wall-normal grid size of the grid cells adjacent to the wall. The zero-gradient condition is prescribed for both *k* and *ω* for open curtains and exhaust fans.

## Footnotes

1 see http://bess.illinois.edu

